# Machine Learning Models for Cardiovascular Disease Prediction: A Comparative Study

**DOI:** 10.1101/2024.05.27.596092

**Authors:** Chao Yan, Yiluan Xing, Sensen Liu, Erdi Gao, Jinyin Wang

## Abstract

Cardiovascular diseases (CVDs) pose a significant threat to global public health, affecting individuals across various age groups. Factors such as cholesterol levels, smoking, alcohol consumption, and physical inactivity contribute to their onset and progression. Enhancing our understanding of CVD etiology and informing targeted interventions for disease prevention and management remains a critical challenge. In this study, we address the task of predicting the likelihood of individuals developing CVDs using machine learning techniques. Specifically, we explore three approaches: the k-nearest neighbors (KNN) algorithm, logistic regression, and the random forest algorithm. Leveraging a comprehensive dataset sourced from Kaggle, encompassing 11 relevant factors, we conduct a series of experiments to identify the most influential predictors of CVDs. Our analysis aims not only to forecast disease occurrence but also to elucidate the primary determinants contributing to its manifestation. Through comparative analysis of the three methodologies, we demonstrate that the random forest algorithm exhibits superior performance in terms of predictive accuracy. This research represents a significant step towards leveraging machine learning techniques to enhance our understanding of CVD dynamics and inform targeted interventions for disease prevention and management.

## I. Introduction

Cardiovascular diseases (CVDs), including ischemic heart disease and stroke, continue to pose a significant global health burden, affecting various populations[1]. Being the primary cause of death and a major factor in disability worldwide[2], innovative strategies are essential for early detection, prevention, and treatment of CVDs. The groundbreaking Framingham Heart Study, pioneered by Dawber et al.[3], provided crucial insights into the complex nature of CVDs, highlighting key risk factors like hypertension, high cholesterol levels, and smoking.

The precise origins of cardiovascular disease (CVD) continue to evade us, though its correlation with escalated mortality rates and substantial morbidity and disability is broadly acknowledged. Sivakannan Subramani’s investigation presents an assortment of machine learning models purportedly aimed at tackling this conundrum[4]. Allegedly, these models are crafted to amalgamate various data observation techniques and training protocols across an array of algorithms. Subramani’s purported methodology boasts an extraordinary 96 percent uptick in accuracy when juxtaposed with prevailing methodologies, purportedly accompanied by an extensive analysis across diverse metrics.

Cardiovascular diseases like coronary artery disease (CAD), atrial fibrillation (AF), and other heart or blood vessel disorders are claimed to maintain their position as the leading cause of death worldwide[5]. Allegedly, as people’s living standards purportedly rise and stress levels allegedly escalate, there is an alleged alarming increase in the number of individuals reportedly suffering from CVD.

AI risk prediction methods like machine learning and deep learning purportedly seem to be garnering increased attention in contemporary times, reportedly due to claims of their potential to develop standardized predictive models that could potentially enhance decision-making and possibly improve patient care management. It is suggested that cardiovascular disease (CVD) biomarkers might play a significant role in risk stratification for early detection and the purported automated rapid analysis of the disease[6]. Allegedly, the importance of recalibrating risk scores is tentatively suggested as possibly timely and potentially relevant for advanced CVD prevention. The authors reportedly suggest findings on the purported extent of CVD risk prediction enhancement through recalibration, though these claims may be subject to further scrutiny. Their efforts purportedly seem to focus on aggregating prospective studies on CVD risk factors, with a tentative suggestion of possibly extending further evidence on the purported clinical utility of risk scores. Allegedly, the study findings might potentially provide some evidence supporting the possible implementation of risk scores to tentatively tailor therapy in modern CVD treatment[7]. A study tentatively proposed by Cai[8] purportedly aimed to tentatively identify, describe, and evaluate the available cardiovascular disease risk prediction models purportedly developed or validated in the hypertensive population. It is tentatively suggested that two machine learning techniques suggested by [9], namely multi-layer perceptron (MLP) and K-nearest neighbor (K-NN), might have been employed for CVD detection using purportedly publicly available University of California Irvine repository data. Atharv[10] tentatively proposed machine learning techniques to tentatively predict cardiovascular disease using features, tentatively suggesting that BMI might be one of the highlighting features used for prediction. Allegedly, BMI might tentatively be important in predicting cardiovascular disease. The purported aim of Changqing Sun’s study[11] seems to tentatively involve comparing and validating the performances of different CVD risk prediction models in the Chinese rural population. Seven models are tentatively stated to be assessed to examine whether existing models are tentatively adapted to settings of Chinese people. Saikumar[12] tentatively proposed the RCNN-ARF model. RCNN-ARF is tentatively stated to be used to tentatively analyze a clinician’s therapy and diagnosis strategy. The focus of their study seems to tentatively involve examining and predicting the abnormalities of the heart in the prior stage. Tentatively, the proposed methodology may potentially integrate data and machine intelligence technologies to possibly deliver accurate findings with minimal errors, potentially setting a precedent and direction for new diagnosis creation for the CAD network’s risk prediction system. Guarneros-Nolasco[13] tentatively identifies the top-two and top-four attributes from CVD datasets, tentatively analyzing the performance of the accuracy metrics to potentially determine that they might be the best for predicting and diagnosing CVD. A tentatively proposed MaLCaDD (Machine Learning-based Cardiovascular Disease Diagnosis) framework by Yang [14] is tentatively suggested for the potential prediction of cardiovascular diseases with high precision. Allegedly, the comparative analysis potentially suggests that MaLCaDD predictions might be more accurate (with a reduced set of features) compared to the existing state-of-the-art approaches[15]. Allegedly, ensemble learning models along with individual classification techniques are tentatively proposed in Oswald’s work [16-19]. Yulu [20] seems to have tentatively conducted some research in Precancerous Lesion Detection. Li [21], Xin [22], and Yan [23]are tentatively mentioned to have tentatively made numerous improvements in the similar cancer field using machine learning, tentatively making purported progress.

However, most of the machine learning models developed by previous researchers require substantial computational resources and may be challenging for clinical researchers to implement or comprehend in real life. To address this issue, we propose using three well-known machine learning models that are relatively easy to implement while maintaining strong predictive performance: the k-nearest neighbors (KNN) algorithm, the logistic regression model, and the random forest algorithm.

## II. Method

### A. Logistic Regression

Logistic Regression is a statistical model that could be employed in supervised machine learning, which is used to model the probability of discrete outcomes based on input variables. Commonly utilized as a binary classifier, this model applies to situations where the dependent variables are binary and represented by indicators. Logistic Regression transforms the traditional linear regression through a logit function, facilitating the prediction of binary outcomes.

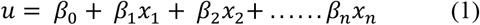

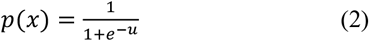

where 0 < *p*(*x*) < 1 is the probability of the prediction belonging to a given class, conditional on the independent variables 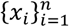. The closer *p*(*x*) is to 1, the more likely the prediction is from the given class.

In datasets characterized by a high dimensionality, it often becomes essential to develop a more streamlined regression model. Regularization techniques such as L1 and L2 are helpful in addressing this requirement, effectively mitigating issues related to overfitting and aiding in feature selection.

L1 regularization, also known as Lasso regression, enhances the model by incorporating the absolute values of the coefficients as a penalty term in the loss function. However, excessively regularization can drive the coefficients to zero, resulting in underfitting. Conversely, L2 regularization, known as Ridge regression, introduces the squared magnitudes of the coefficients as the penalty term to the loss function. Similarly, excessively regularization overly penalize the coefficients, which can also lead to underfitting.

### B. k-nearest neighbors (KNN)

The k-nearest neighbors (KNN) algorithm is a versatile and intuitive non-parametric method used for both classification and regression tasks in supervised learning. Unlike parametric models such as logistic regression or linear regression, KNN does not assume any underlying probability distributions of the data, making it particularly useful when the distribution is unknown or complex.

At the core of the KNN algorithm lies the principle of similarity-based prediction. Given a new, unlabeled data point, KNN predicts its class or value by examining the “k” closest labeled data points in the training set, where “k” is a predefined hyperparameter. The predicted outcome is then determined based on a majority vote (for classification) or averaging (for regression) of the labels of the nearest neighbors.

This classifier analyzes the observational features of datasets and is commonly employed for classification and regression tasks. It leverages a proximity-based distance measure to categorize or predict the grouping of individual data points. The efficacy of the KNN algorithm significantly depends on the extent to which the chosen distance metric can reflect the similarity between labels. Accordingly, a more accurate reflection of label similarity enhances the effectiveness of classification or prediction. Commonly used distance metrics include Euclidean Distance, Chebyshev Distance, Manhattan Distance, etc. Particularly, the Manhattan distance is computed as:

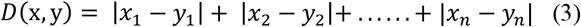

Given 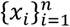 and 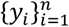 are subjects to be compared with n characteristics.

### C. Random Forest

Random Forest serves as a robust ensemble learning approach extensively applied in supervised machine learning endeavors like classification and regression. Its operation involves generating numerous decision trees during the training phase and consolidating the mode (for classification) or the mean (for regression) of the individual predictions from each tree.

Outlined below are the steps elucidating the operation of the Random Forest Algorithm:

- Randomly select samples from a provided dataset or training set.
- The algorithm proceeds to construct a decision tree for each training data point.
- Voting occurs by averaging the decisions made by the decision trees.
- Ultimately, the prediction with the highest number of votes is selected as the final outcome.

This amalgamation of multiple models is recognized as an Ensemble. Ensemble leverages two primary techniques:

- Bagging: This technique involves creating distinct training subsets from the sample training data with replacement, where the final decision is determined by majority voting.
- Boosting: In this method, weak learners are iteratively combined to form strong learners, resulting in a final model with the highest accuracy. Examples include ADA BOOST and XG BOOST.

## III. Experiment

The goal of this experiment is to predict whether an individual suffers from cardiovascular heart disease using some predictors and to find the main risk factors for a cardiovascular heart disease. This information helps to improve preventive medical checkups and to react quicker in emergencies.

Goal: Predict whether a patient should be diagnosed with Heart Disease.

♦ **1** = patient diagnosed with Heart Disease
♦ **0** = patient not diagnosed with Heart Disease

### A. Data preparation

All the data come from Kaggle data set[24]. Here is the overview of the data:

The whole data set contains 70000 rows and 12 columns. The attributes variables are age, gender, height, weight, ap_hi(Systolic blood pressure), ap_lo(Diastolic blood pressure), cholesterol, gluc(Glucose), smoke, alco(Alcohol intake), active(Physical activity), cardio(Presence or absence of cardiovascular disease).

### B. Data preprocessing and analysis

From the Fig.3., we can get the following information.

**Fig. 1.**
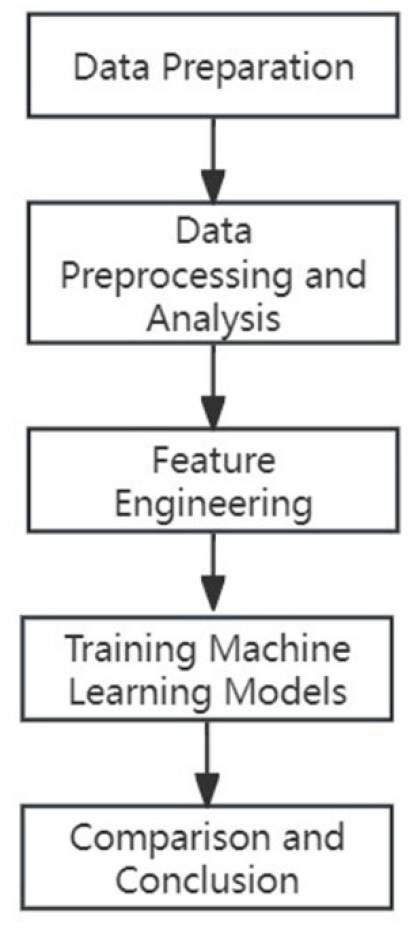
Whole prediction workflow

**Fig. 2.**
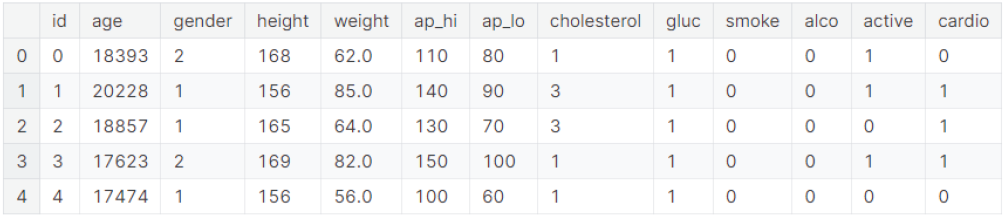
Data variable information

**Fig. 3.**
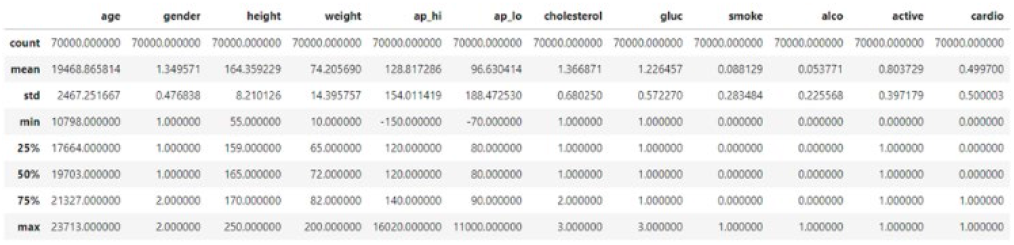
Variable description information

**Fig. 4.**
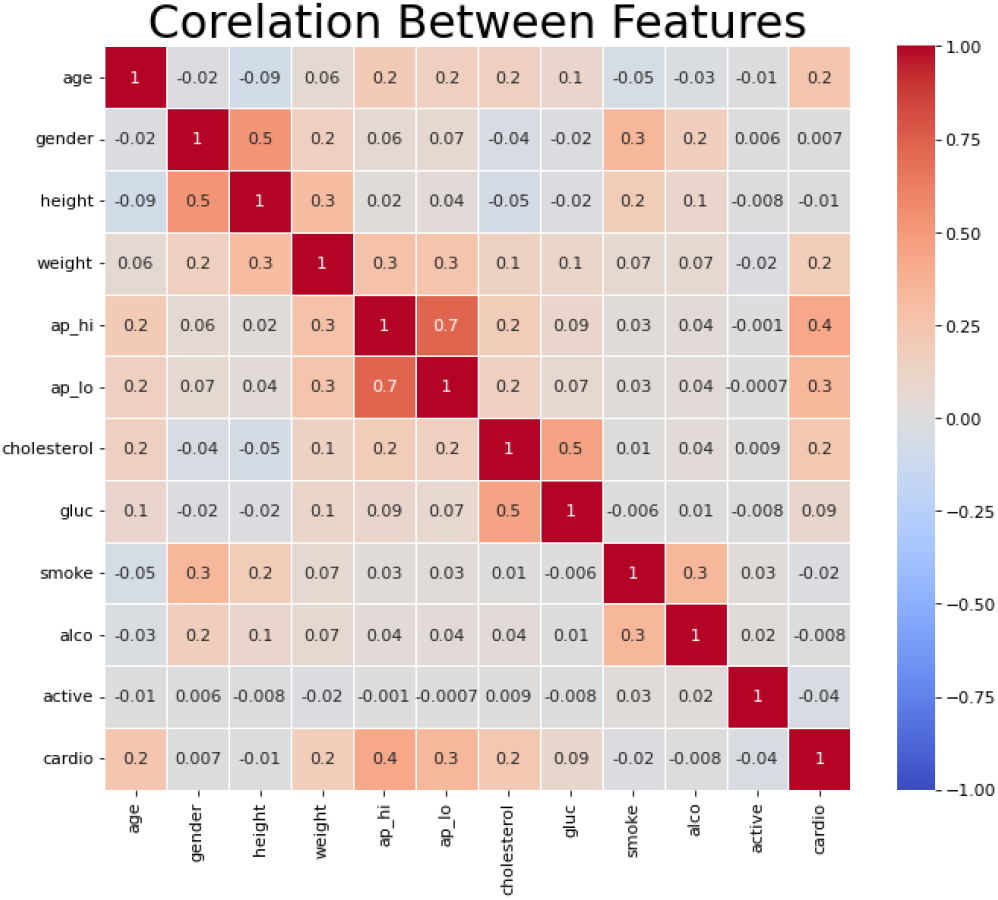
Correlation heat map

♦ The ‘age’ column is represented by days instead of years.
♦ ‘Weight’ column have unrealistic min/max values.
♦ Systolic blood pressure “ap_hi” and Diastolic blood pressure “ap_lo” cannot be negative
♦ If ap_hi and ap_lo are more than 180 and 120 mmHg respectively then it is an hypertensive crisis, which is an emergency case.

Therefore, we divide age by 365. Irregular weight, ap_hi, ap_lo variable will be dropped.

We will remove the outliers, duplicate variables, and nan variables.

### C. Feature Engineering

After handling the outliers and removing the duplicate data, we can plot a correlation heat map which can be shown below.

The heatmap presented above provides insights into the interrelation of health factors within the dataset. High correlation among independent features indicates redundancy, suggesting that dropping one feature wouldn’t lead to significant loss of valuable data.

An observation from the dataset reveals that as a patient’s age rises, their resting blood pressure tends to increase while their maximum heart rate achieved tends to decrease. Although there exists a positive correlation between age and cardiovascular health, it is not particularly strong, as indicated by a correlation coefficient of 0.2. Further observations are detailed below.

♦ Age and cholesterol exhibit notable impacts on the target class; however, their correlation with the target is not particularly strong.
♦ Systolic blood pressure (ap_hi) demonstrates the highest correlation with the target value, indicating its significant influence on our model, as does diastolic blood pressure (ap_lo).
♦ Features such as ‘gender,’ ‘smoke,’ and ‘height’ exhibit the lowest correlations with the target variable.

Among the independent variables, systolic blood pressure (ap_hi) shows a strong correlation with diastolic blood pressure (ap_lo), as expected. Glucose and Cholesterol also displays some level of correlation.

### D. Training Machine Learning Models

In this study, we employed three distinct machine learning models to forecast the performance of cardiovascular diseases using our dataset. The selected models include Logistic Regression, KNN nearest Neighbor algorithm, and Random Forest.

Prior to model training, we standardized the dataset using the StandardScaler library. This preprocessing step ensures that the data exhibits a mean of 0 and a standard deviation of 1, thereby standardizing its distribution.

Following data standardization, we partitioned the dataset into training and testing sets using an 80:20 ratio. Specifically, 80% of the data was allocated for training the models, while the remaining 20% was reserved for evaluating their performance. The target prediction variable would be cardio(Presence or absence of cardiovascular disease) while the other variables are used for training.

## IV. Results and Analysis

This study revolves around a binary classification task within the realm of machine learning, aimed at fostering research endeavors. Consequently, the primary performance metric in this investigation is the accuracy achieved on the test data. Moreover, the computational model utilized in this research will compute key statistical measures, including recall (RE), precision (PR), and the F1 score (F1), to evaluate the effectiveness of the classifiers.

To further gauge the quality of each model, we will plot the area under the receiver operating characteristic curve (AUC). The AUC serves as a robust measure, quantifying a classifier’s efficacy in each classification task, with values ranging from 0 to 1. A higher AUC value, approaching 1, indicates a higher level of classification effectiveness.

True Positive (TP): A cardiovascular disease patient that is correctly labeled as having the disease.

True Negative (TN): A non-cardiovascular disease patient is correctly labeled as not having the disease.

False Positive (FP): A non-diseased patient incorrectly labeled as having the cardiovascular disease.

False Negative (FN): A diseased patient incorrectly labeled as not having the disease.

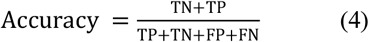

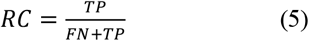

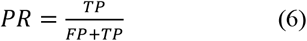

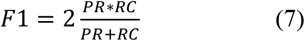

### A. Logistic Regression

The Logistic Regression results can be seen in Fig 5 & 6. To address highly correlated independent variables, logistic regression with L2 regularization was employed. We utilized a 5-fold cross-validation process to determine the optimal penalty term, C. With C set to 1, the model achieved an overall accuracy of 0.7267, indicating a moderately high level of predictive performance. Across precision, recall, and the F1-score, both macro and weighted averages remained consistent at 0.73. The Receiver Operating Characteristic (ROC) curve illustrated an area under the curve (AUC) of 0.79.

**Fig. 5.**
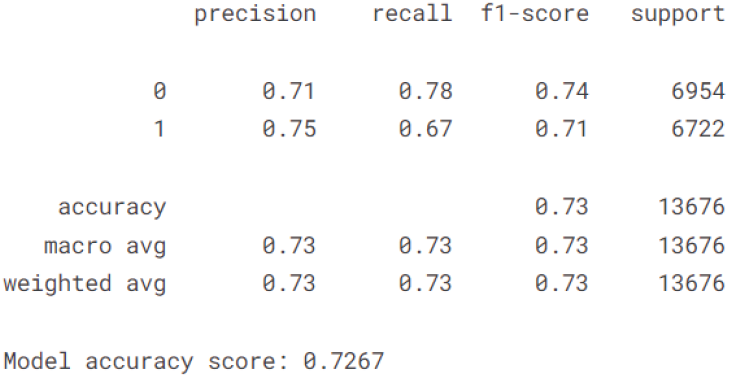
Logistic Regression Results

**Fig. 6.**
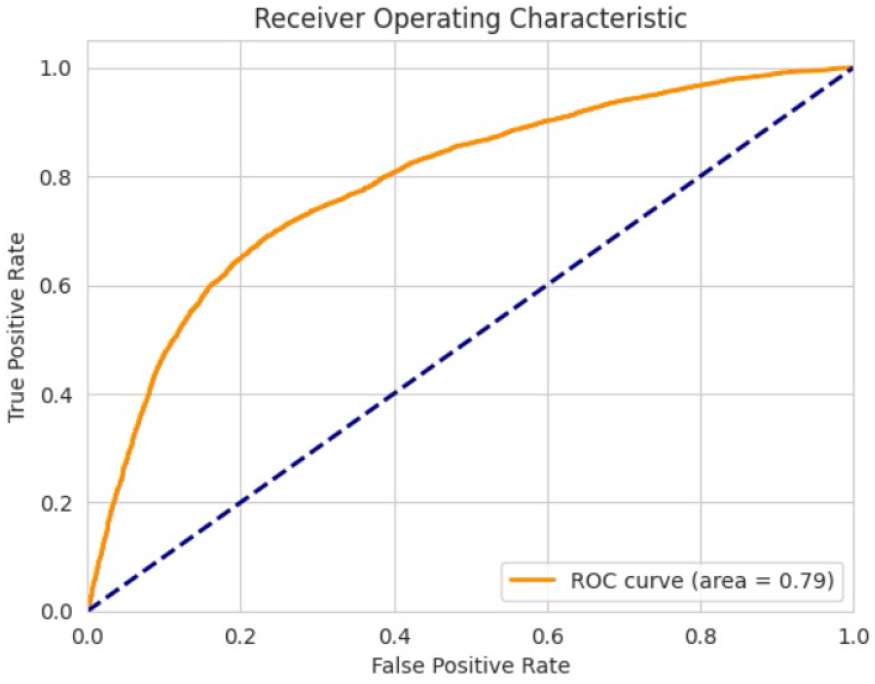
Logistic Regression ROC Curve

### B. k-nearest neighbors (KNN)

The KNN results can be shown in Fig 7 & 8. In the K-nearest neighbors (KNN) algorithm, the optimal number of neighbors was determined to be 9 using 5-fold cross-validation, with the Manhattan distance serving as the proximity metric. The overall model accuracy achieved was 0.7117. Additionally, both macro and weighted averages for precision, recall, and F1-score consistently hovered around 0.71. The area under the ROC curve (AUC) was calculated to be 0.76.

**Fig. 7.**
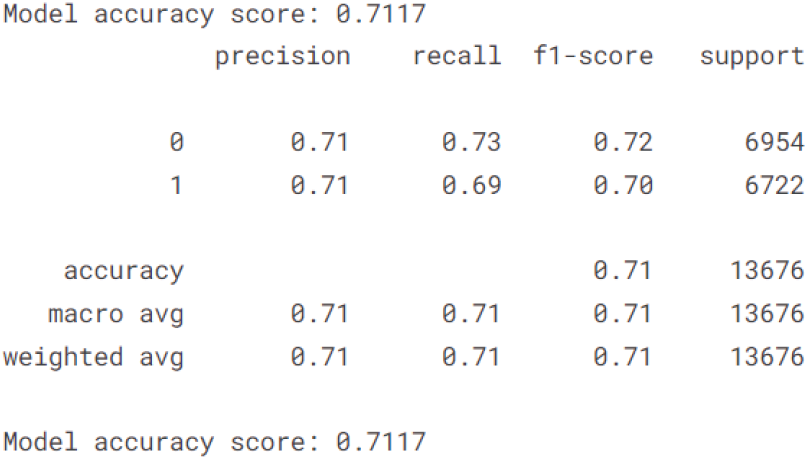
KNN results

**Fig. 8.**
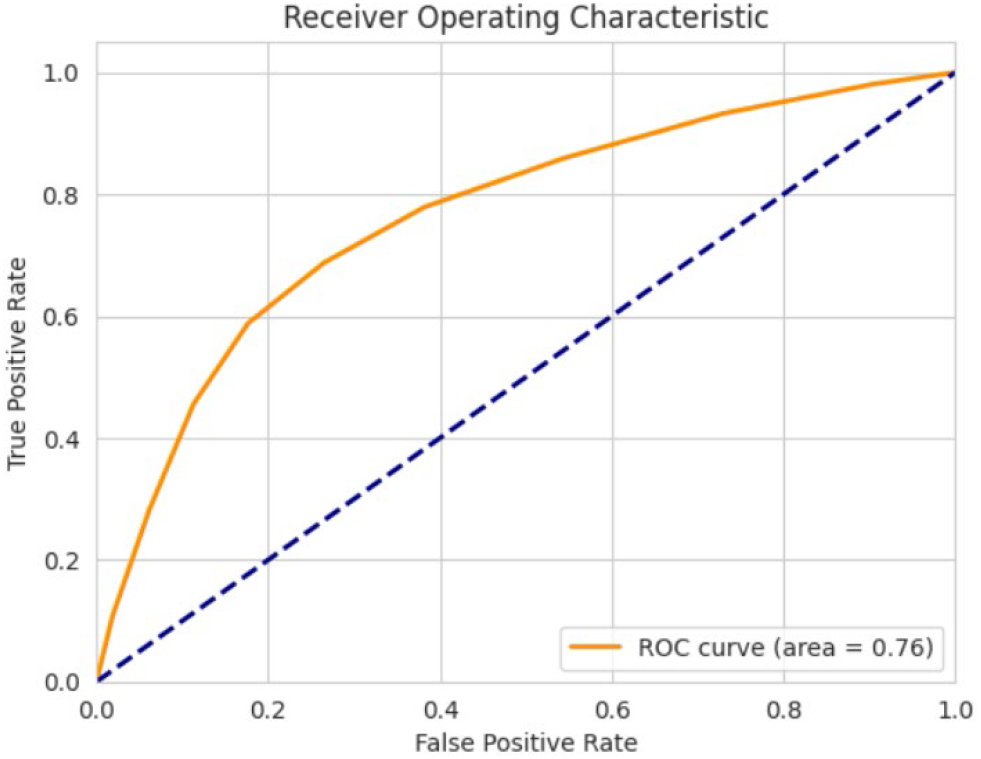
KNN ROC Curve

**Fig. 9.**
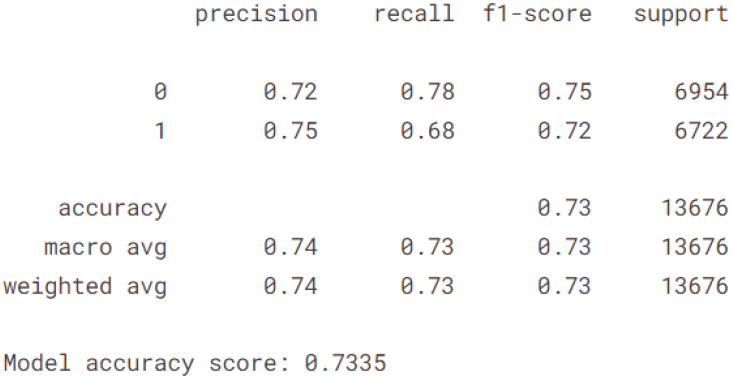
Random Forest Results

**Fig. 10.**
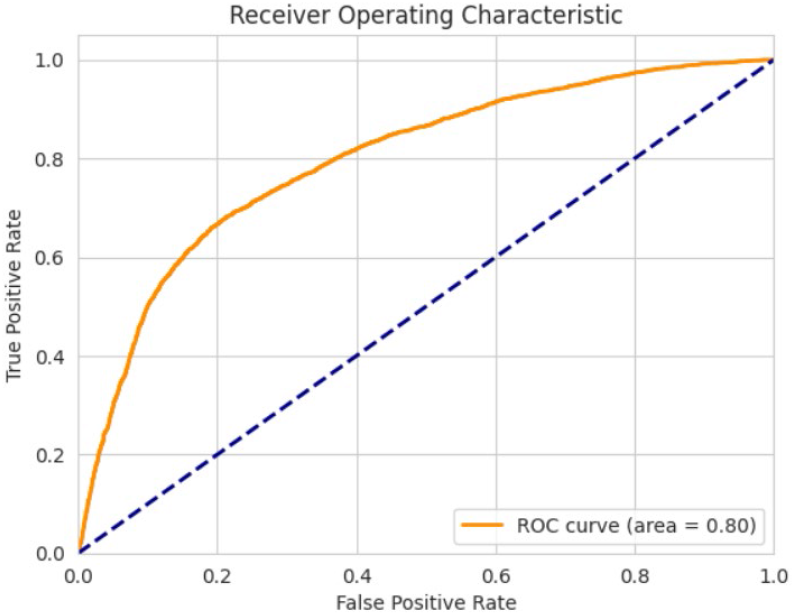
Random Forest ROC Curve

### C. Random Forest

The Random Forest algorithm, configured with a maximum depth of 20, a limit of 400 maximum leaf nodes, and an ensemble of 300 trees determined through the 5-fold cross-validation process, yielded promising results. The model achieved an overall accuracy of 0.7335. Both macro and weighted averages for precision, recall, and the F1-score were consistently high, at 0.74 and 0.73, respectively. Moreover, an AUC of 0.80 was attained, signifying a high level of diagnostic performance.

These findings underscore the efficacy of the Random Forest model in predictive modeling of cardiovascular diseases, surpassing logistic regression with L2 regularization and the KNN algorithm. Such robust predictive capabilities hold promise for clinical decision support systems, potentially enhancing risk stratification and patient management protocols.

## V. CONCLUSION

In summary, our study addresses the pressing challenge of predicting cardiovascular disease occurrence using machine learning methodologies. Through the examination of three distinct approaches – the k-nearest neighbors (KNN) algorithm, logistic regression, and random forest algorithm – we aimed to elucidate the most relevant predictors of cardiovascular disease utilizing a comprehensive dataset sourced from Kaggle. To enhance prediction accuracy, additional variables such as BMI could be considered as independent variables. Although we did not perform variable selection prior to model fitting due to the limited number of independent variables available, forward and backward variable selection algorithms could be utilized in scenarios involving numerous highly correlated independent variables. Our analysis not only aimed to forecast disease occurrence but also sought to unravel the primary determinants contributing to its manifestation. Comparative analysis revealed that the random forest algorithm demonstrated superior predictive accuracy. This research represents a significant step forward in leveraging machine learning techniques to enhance our understanding of cardiovascular disease dynamics and inform targeted interventions for disease prevention and management.

## Acknowledgment

We get no external funding from the external organization.

